# Local enrichment of cardiolipin to transient membrane undulations

**DOI:** 10.1101/2025.01.17.633669

**Authors:** Christopher T. Lee, Kailash Venkatraman, Itay Budin, Padmini Rangamani

## Abstract

Organelles such as mitochondria have characteristic shapes that are critical to their function. Recent efforts have revealed that the curvature contributions of individual lipid species can be a factor in the generation of membrane shape in these organelles. Inspired by lipidomics data from yeast mitochondrial membranes, we used Martini coarse-grained molecular dynamics simulations to investigate how lipid composition facilitates membrane shaping. We found that increasing lipid saturation increases bending rigidity while reducing the monolayer spontaneous curvature. We also found that systems containing cardiolipin exhibited decreased bending rigidity and increased spontaneous curvature when compared to bilayers containing its precursor, phosphatidylglycerol. This finding contradicts some prior experimental results that suggest that bilayers containing tetraoleoyl cardiolipin have greater rigidity than dioleoyl phosphatidylcholine bilayers. To investigate this discrepancy, we analyzed our simulations for correlations between lipid localization and local curvature. We found that there are transient correlations between curved lipids such as cardiolipin (CDL) and phosphatidylethanolamine (PE) and curvature; these interactions enrich specific bilayer undulatory modes and cause bilayer softening. Furthermore, we show that curvature-localization of some lipids such as cardiolipin can influence lipids in the opposing leaflet. These observations add to the emerging evidence that lipid geometric features give rise to local interactions, which can cause membrane compositional heterogeneities. The cross-talk between composition-driven tuning of membrane properties and membrane shape has implications for membrane organization and its related functions.

**SIGNIFICANCE:** The material properties of phospholipid membranes are a function of the lipid composition. The abundance of cardiolipin in the mitochondrial inner membrane implies a functional role for this special lipid. We explore the interactions of cardiolipin with other lipids with varying lipid saturation using coarse-grained molecular dynamics simulations. We find that membranes containing cardiolipin have higher spontaneous curvature and lower bending rigidity when compared to membranes without cardiolipin – in line with prior models and experiments. We also show that the low bending rigidity of cardiolipin-containing symmetric, and flat bilayered systems is due to the transient partitioning of cardiolipin to undulations due to curvature sensing. This is a mechanism for forming lateral membrane heterogeneities in otherwise symmetric systems.

## INTRODUCTION

Mitochondria are central to energy production in cells; their unique and characteristic ultrastructure is fundamental to their function and performance (1, 2). Unlike most other organelles, mitochondria have two phospholipid bilayers called the outer mitochondrial membrane (OMM) and inner mitochondrial membrane (IMM). The IMM, where the machinery for the electron transport chain and adenosine triphosphate (ATP) synthesis are localized, has a highly curved, folded, and tubulated shape (3, 4). This folded geometry with a high surface area to volume ratio is critical for energy production (1, 2, 5).

Recently, many studies have pointed to the important role of mitochondrial morphology in maintaining its physiological function (6). The loss of the characteristic IMM ultrastructure is associated with defects in energy production and is observed in many diseases (7, 8). Recently, we explored the consequence of changing lipid saturation on the structure and function of yeast mitochondria (9). Using a promoter strategy to control the expression of the sole lipid desaturase in yeast, we were able to tune the extent of lipid saturation in the mitochondrial membranes. We found that increasing lipid saturation led to an abrupt loss of IMM ultrastructure; that is, there is a ‘breaking point’ of lipid saturation which, when exceeded, leads to loss of cristae folds. This loss of cristae is driven in part by the dissociation ATP synthase dimers (9). We further observed that knocking out cardiolipin synthase (*CRD1*), which causes the accumulation of phosphatidylglycerol (PG) instead of CDL, shifts this ‘breaking point’ such that a lesser increase in saturation leads to loss of shape (notably without shifting ATP synthase oligomerization). Previously, we used continuum modeling to explain this abrupt loss of cristae structure as a snapthrough instability (9, 10). However, such continuum models depend on the input material properties such as bending modulus and do not account for the local lipid composition or lipid-lipid interactions. Briefly, the energy of a membrane bent into a particular shape is often modeled using the Helfrich-Canham (HC) Hamiltonian (11–13),

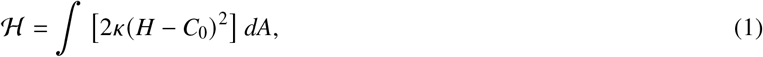

where *κ* is the bending modulus, *H* is the mean curvature, and *C*_0_ is the spontaneous (*mean*) curvature. In the HC theory, there are two material parameters *κ* and *C*_0_ which dictate energy required to bend a membrane. These parameters are also sophisticated functions of the lipid chemistry and composition. They can be estimated from molecular dynamics (MD) simulations (14–19) where advances in lipid force field parameterizations enable the study of complex lipid mixtures (20), Fig. 1. Analyses of mean height fluctuations of the bilayer enable the characterization of the bending modulus, *κ*; while the first moment of the lateral pressure profile is related to the bending moment, or the product of the bending modulus and spontaneous curvature, *KC*_0_ (Fig. 1B-C).

**Figure 1:**
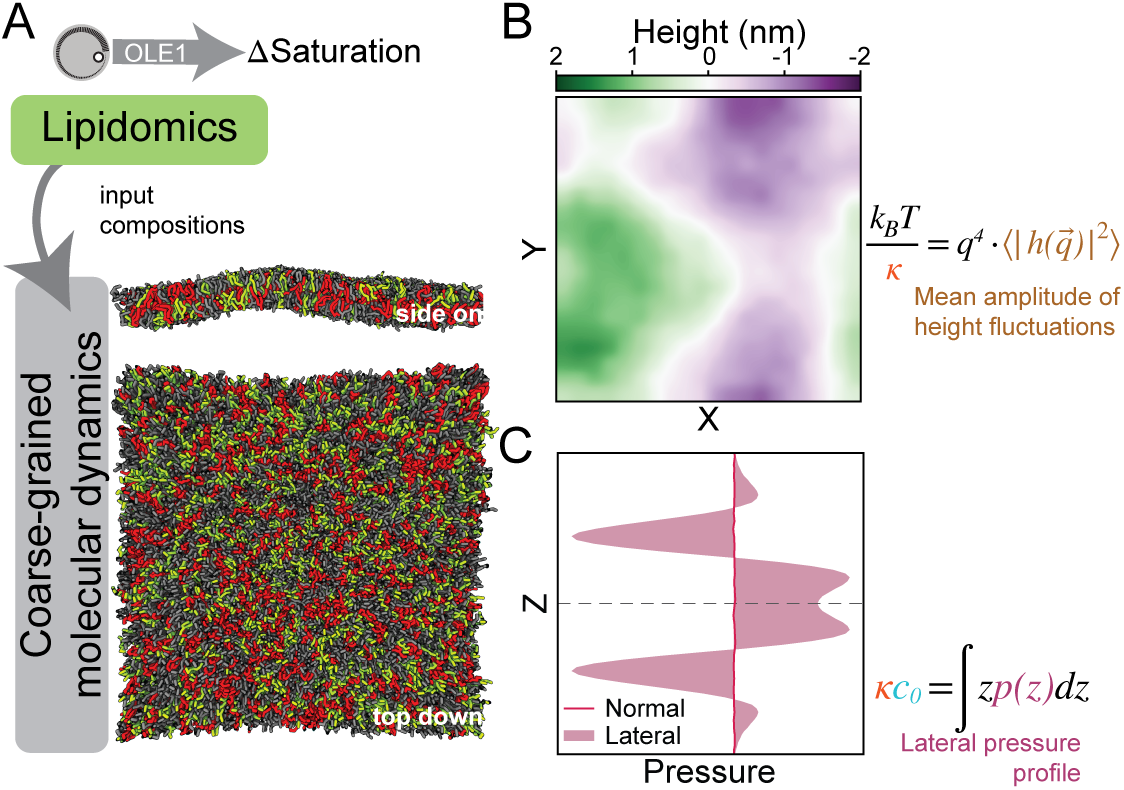
Overview of the approach connecting coarse-grained molecular dynamics simulations to continuum modeling. A) Lipid compositions from lipidomics of yeast mitochondria with the extent of lipid saturation under experimental control are input into coarse-grained molecular dynamics simulations. B) Analysis of mean height fluctuations of the bilayer enables the characterization of the bending modulus, *κ*. C) The first moment of the lateral pressure profile is related to the bending moment, or the product of the bending modulus, *κ* and spontaneous mean curvature, *c*_0_.

In this work, we seek to answer the following questions: How does the presence of cardiolipin affect the mechanical properties lipid bilayers? How does the presence or absence of cardiolipin shift the point of lipid saturation that breaks cristae curvature formation? To answer these questions, we used coarse-grained molecular dynamics simulations (CGMD) with experimentally informed lipid compositions.

The prior literature, consisting of both experimental and computational approaches, presents conflicting perspectives on the effect of CDL on the bilayer. On one hand, experimental efforts investigating the partition of CDL to membrane tubes pulled from giant unilamellar vesicles suggest that it has a conical shape, which imparts it with a large spontaneous curvature (21). Similarly, quantification of extracts of mitochondrial lipids also suggest that CDL rich membranes have a very low bending rigidity (22). In contrast, measurements from scattering experiments suggest that CDL has a weaker negative spontaneous curvature (cf. Fig. S1) than PE (23–25) and an increased rigidity compared to 1,2-dioleoyl-sn-glycero-3-phosphocholine (DOPC) (26). Investigation using coarse-grained molecular dynamics (CGMD) suggests a reduced bending rigidity (27, 28), and a preference for CDL to partition to regions of negative curvature (29–32). Reports from simulations of atomistic systems report larger bending rigidities of CDL than 1-palmitoyl-2-oleoyl-sn-glycero-3-phosphocholine (POPC) (33, 34) and a weaker absolute spontaneous curvature for CDL than 1-palmitoyl-2-oleoyl-sn-glycero-3-phosphoethanolamine (POPE) (34).

In this work, we aim to reconcile these conflicting perspectives by investigating the mechanical properties of membranes with experimentally-informed lipid compositions. By modeling membranes with lipid compositions informed by lipidomics of yeast mitochondria, we predict how the compositional changes influence the material properties of the membrane, Figs. 1 and 2. We find that increasing lipid saturation leads to an increase in the bending rigidity of the membrane and a general decrease in spontaneous curvature. Our CGMD simulations also revealed that the CDL containing systems have a diminished bending rigidity compared with PG containing systems. While these findings are in general agreement with prior CGMD simulation studies (27), our estimated bending moduli are lower and spontaneous curvature larger than expected from experiments of CDL containing systems (23–26). Examination of the height fluctuation spectra of CDL containing systems suggests that these systems exhibit spectral features deviating from the predictions of the HC model, Fig. 3C.

**Figure 2:**
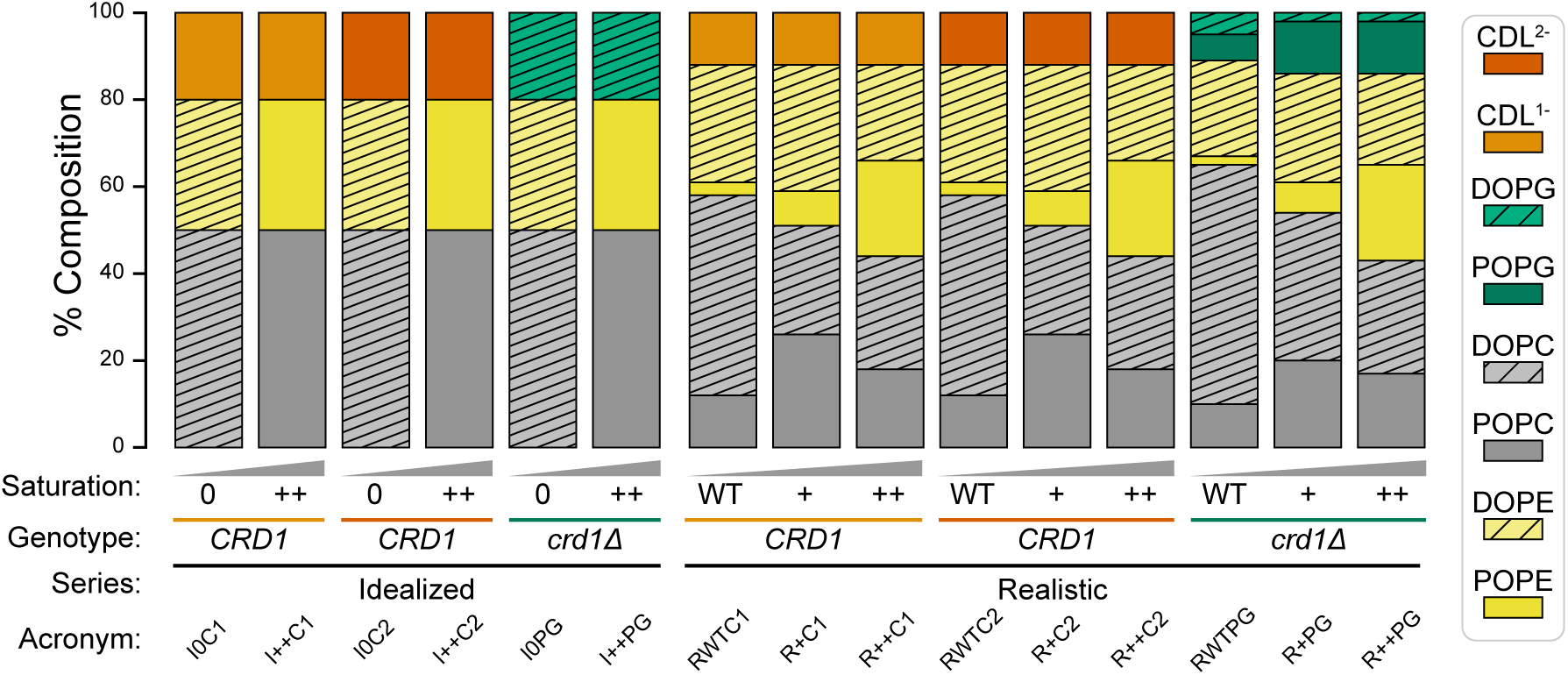
Compositions of the modeled systems. The systems are separated into two cohorts: idealized systems (left hand side) which are composed of either dioleoyl (DO) or palmitoyl (PO) forms of phosphatidylcholine (PC) and PE lipids and realistic compositions (right hand side) based on lipidomics data of yeast mitochondria (9). Systems were further categorized by the presence of CDL^1 –^, CDL^2 –^, or PG. The CDL modeled is tetraoleoyl cardiolipin (TOCL) and further differentiated by charge state. Systems containing PG correspond to cardiolipin synthase knock out (*crd1Δ*) yeast mutants. Systems will be referred to by the following nomenclature: ({R,I}{WT/0, +, ++}{C1, C2, PG}) (e.g., RWTC1 corresponds to the realistic CDL^1 –^ wild type condition, and I++PG the idealized PG system at elevated saturation).

**Figure 3:**
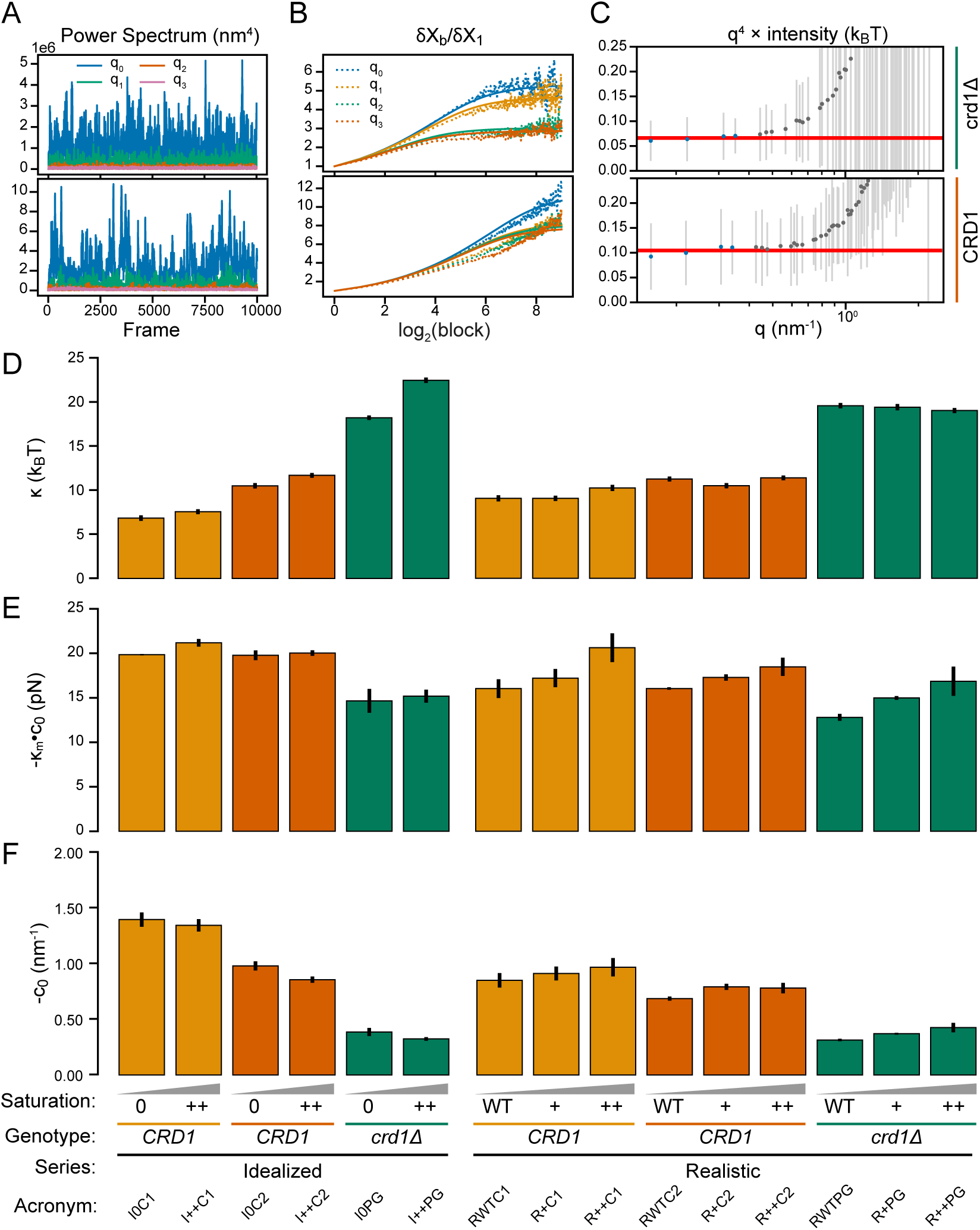
Quantification of bending modulus, bending moment, and spontaneous curvature for each system. For panels A, B, and C, the top panels are derived from system I0PG and bottom from I0C2. A) An example power spectrum (i.e., the first *i* undulatory modes, *q*_*i*_), B) The extent of correlation within the power time series for each undulatory mode is estimated by fitting to the block average. C) Plot of the height power spectra showing the linear fit at large wavenumber regimes. The mean values for each undulatory mode are shown by a dot and the estimated standard deviation shown by gray lines. D) Estimated bending moduli for all systems. Error bars correspond to the standard deviation derived from parametric bootstrapping of data in C. E) The bending moment is calculated as a product of monolayer bending rigidity and monolayer spontaneous curvature. F) The monolayer spontaneous curvature for each system derived by substituting the estimated bending rigidity and bending moment in D and E.

Considering that the unique spectral features of CDL containing systems could be caused by heterogeneities in membrane organization, we investigated the distributions of lipids around a given lipid species and found that there are preferential interactions. Investigating this further, beyond lipid chemical interactions, we found that there are transient correlations of lipids even with the curvature of the membrane. We observed that CDL and PE both exhibit a preference to localize with negative mean curvature, but only CDL exhibits depletion across the bilayer. Altogether, the kinetics of curvature-generating or -stabilizing interactions between lipids can lead to bilayer softening, which we propose as the cause of the deviations from the HC model (35).

## METHODS

### Molecular dynamics simulations

Symmetric bilayers with compositions derived from systems described previously in Venkatraman et al. [9] (also shown in Figs. 2 and S2 and Table S1), were modeled by coarse-grained molecular dynamics using the Martini 2.2 force field parameters (36). Systems of approximately 40 × 40 × 25 nm and 15 × 15 × 25 nm (referred to as the small systems) were constructed using insane.py (37). All systems were solvated with water and charge neutralized by 150 mM NaCl using the workflow provided by insane.py. The small systems were solvated with 90 % water and 10 % antifreeze molecules to prevent freezing of the system. Minimization and equilibration protocols were adapted from CHARMM-GUI Martini Maker (38) and summarized in the Section B.1. Production runs extend for a total of 5 µs of simulation time from the equilibrated structures with a 20 fs timestep. Small systems were simulated for a total of 3 µs followed by 200 ns of simulation with a 5 ps write out frequency for calculation of system stresses. Notably the simulation lengths reported reflect the actual simulation length without factoring in the conventional *post hoc* time scaling of Martini simulations.

The pressure was maintained in production simulations using the Parinello-Rahman Semiisotropic barostat with *τ*_*p*_ of 12 ps, compressibility 3 × 10^−4^ bar^−1^, at 1 bar pressure (39). For all simulations, electrostatics were treated using the reaction-field method with a cutoff of 1.1 nm and _*a*_ 15 (40), and the temperature was held at 303 K using the stochastic velocity-rescaling thermostat with *τ*_*t*_ set to 1.0 ps (41).

### Analysis

From the large-box simulations, the bending modulus was estimated by analysis of the height fluctuation spectra, see Section B.1.1. Meanwhile, from the small-box simulations, the spontaneous curvatures were estimated from the moments of the lateral pressure profile and substituting the estimated bending modulus for each system, see Section B.1.3 for detailed approach. Statistical errors are propagated using conventional formulae.

## RESULTS

### Model development

In this work, we focused on lipid compositions inspired by the experimental manipulations presented in Ref. (9) with the goal of understanding how mitochondrial lipid composition could affect global and local properties of the membranes. We simulated these membrane compositions using Martini 2.2 CGMD; the compositions are illustrated in Figs. 2 and S2 with details given in Table S1. To study the consequence of lipid saturation on the membrane, we designed idealized systems composed of either DO or PO forms of PC and PE lipids. These idealized systems establish a comparison for the effect of lipid saturation on bilayer properties. We also designed a ‘realistic’ series of compositions informed by lipidomics. In these series, the population of PC and PE lipids along with lipid tail identities were drawn from lipidomics of mutant yeast (9). As a further manipulation, to investigate the importance of CDL, we compared systems with CDL against systems with PG which correspond to *CRD1* and *crd1Δ* genetic conditions respectively. Within the *CRD1* systems, we further differentiated by the charge of the CDL species. This variation is designed to capture the complex chemical environment around the mitochondrial inner membrane and the fact that the protonation state of CDL within the IMM may vary between the singly, CDL^1 –^, and doubly, CDL^2 –^, deprotonated charge states (42, 43).

Overall, the systems are categorized by three factors: their relative realism (idealistic, realistic), the lipid saturation (WT/0, +, and ++), and the genotype (*CRD1, crd1Δ*). We will use the following abbreviation scheme to refer to each system: {I,R}{WT/0, +, ++}{C1, C2, PG}; the system acronyms are elaborated in Table S1. In addition to these 15 systems, we also modeled symmetric bilayers mimicking the outer and inner leaflet compositions of the IMM with either CDL^1 –^ or CDL^2 –^ hence referred to as OutC1, InnC1, OutC2, and InnC2. Finally, as controls, we also modeled bilayers consisting of only CDL^1 –^ or CDL^2 –^ (referred to as C1-only and C2-only respectively). For each condition, we constructed large (∼40 nm) and small systems (∼15 nm) for height-undulation and lateral pressure analysis respectively.

### Membrane properties as a function of membrane composition

To establish the relationship between the membrane composition and membrane properties, we calculated the bending modulus, bending moment, and the resulting spontaneous curvature based on standard methods (14, 16, 44). For each system, we estimated the bending modulus (Fig. 3D and Fig. S5) by analyzing the height fluctuation spectra (Fig. 3A-C). The reported bending moduli for affected systems are scaled by a constant 20 % to account for systematic biases discovered in the Gromacs simulation engine (45), Fig. S3. The uncertainties of the reported bending moduli are the result of parametric bootstrapping of the data (46). Within the idealized series, we found that increasing lipid saturation leads to an increase in bending rigidity (Fig. 3D). This increase is consistent with prior experimental measurements (47). The trends in the realistic series exhibited modest increases in bending rigidity for the CDL^1 –^ containing systems while CDL^2 –^ and PG containing systems bucked the trend. Consistently across the different conditions, lipid membranes containing CDL had reduced bending rigidity compared to the PG containing systems; systems containing CDL^1 –^ had reduced bending rigidity compared to those with CDL^2 –^. The outer leaflet of the IMM was also found to be less stiff than the inner leaflet, Fig. S5. Consistent with prior literature (27), the bending rigidity of systems with C1-only was (5.8 ± 0.2) k_B_T while C2-only was (11.5 ± 0.2) k_B_T, Fig. S5. Notably, for all CDL containing systems, the power spectrum exhibited spectral features at long wavelength modes, deviating from the constant value predicted by HC, compare top and bottom of Fig. 3B and C.

We also considered how the changes to lipid composition could contribute to the preferred curvatures of the bilayer. We computed the monolayer bending moment, *κ*_*m*_*c*_0_, by numerically integrating the lateral pressure profile of each system, Fig. 3E, Fig. S6. Given that the modeled systems have symmetric compositions between leaflets, we assume that the monolayer rigidity is one-half of the bilayer rigidity, *K*_*m*_ = *K* /2. By substituting the previously estimated bending rigidities into the bending moments, we obtained the monolayer spontaneous curvature, *c*_0_, for each system, Fig. 3F and Fig. S7. We note that the orientation of each leaflet, given by the normal, is chosen to be inward pointing, extending in the same direction as the vector pointing from lipid head groups towards the center of the bilayer. This convention is illustrated in Fig. S1 and implies a positive mean curvature for convex shapes (e.g., a spherical micelle or outer leaflet of a vesicle would have a positive mean curvature) and a negative mean curvature for concave shapes. We found that for the idealized compositions, with increasing saturation, there is a corresponding decrease in spontaneous curvature; for the realistic compositions, the trend is reversed. Across both idealized and realistic compositions, CDL^1 –^ containing systems had the greatest spontaneous curvature, followed by CDL^2 –^, and then PG. The outer leaflet of the IMM with greater CDL content has greater spontaneous curvature than the inner leaflet, Fig. S7. Systems with only CDL^1 –^ exhibited a large spontaneous curvature (−2.8 ± 0.1) nm^−1^; meanwhile only CDL^2 –^ had a weaker spontaneous curvature (−1.52 ± 0.03) nm^−1^. This is in contrast with scattering experiments that report that CDL has a lower spontaneous curvature than PE (23–25).

### Aberrant features in height fluctuation spectra of membranes with cardiolipin

At this point, we were surprised by the low bending rigidities predicted for the CDL containing systems. Moreover, owing to the nature of deriving the spontaneous curvature from the bending moment, the estimated bending rigidity has a strong influence on the predicted spontaneous curvature. Inspection of the approach to quantify statistical inefficiency of the power spectra revealed that the fits to theory were poor for systems with CDL, Fig. 3A, B–bottom. Further plots of the height undulation spectra of CDL containing systems appear to have additional spectral features at long wavelengths. This is a substantial deviation from the HC theory, which predicts a linear dependence of the height undulation spectrum with the fourth power of the wavenumber, Fig. 3C–bottom.

Considering the aberrant spectral features and high spontaneous curvature of CDL containing systems, we considered whether interactions of lipid species could give rise to high curvature complexes (cf. Ref. (35)). To evaluate the propensity of lipid clustering, we compared the mean lipid composition within 1.5 nm of any lipid type against the composition of the bulk system, Fig. 4. We found that certain lipids are enriched or depleted against random from bulk, suggesting that there are heterogeneities driven by the interactions between lipid types. PG lipids are found to be enriched against random with respect to all other lipid types, Fig. 4E, F. CDL on the other hand is found to be depleted against other CDL; CDL^2 –^ around CDL^2 –^ is depleted to a larger extent than CDL^1 –^ around CDL^1 –^ lipids, compare Fig. 4A, B with C, D. Curved lipids CDL and PE were found to be mutually enriched in the local neighborhood, Fig. 4A-D. These observations suggested that lipids are heterogeneously dispersed in these complex compositions, which could lead to emergent mechanical properties.

**Figure 4:**
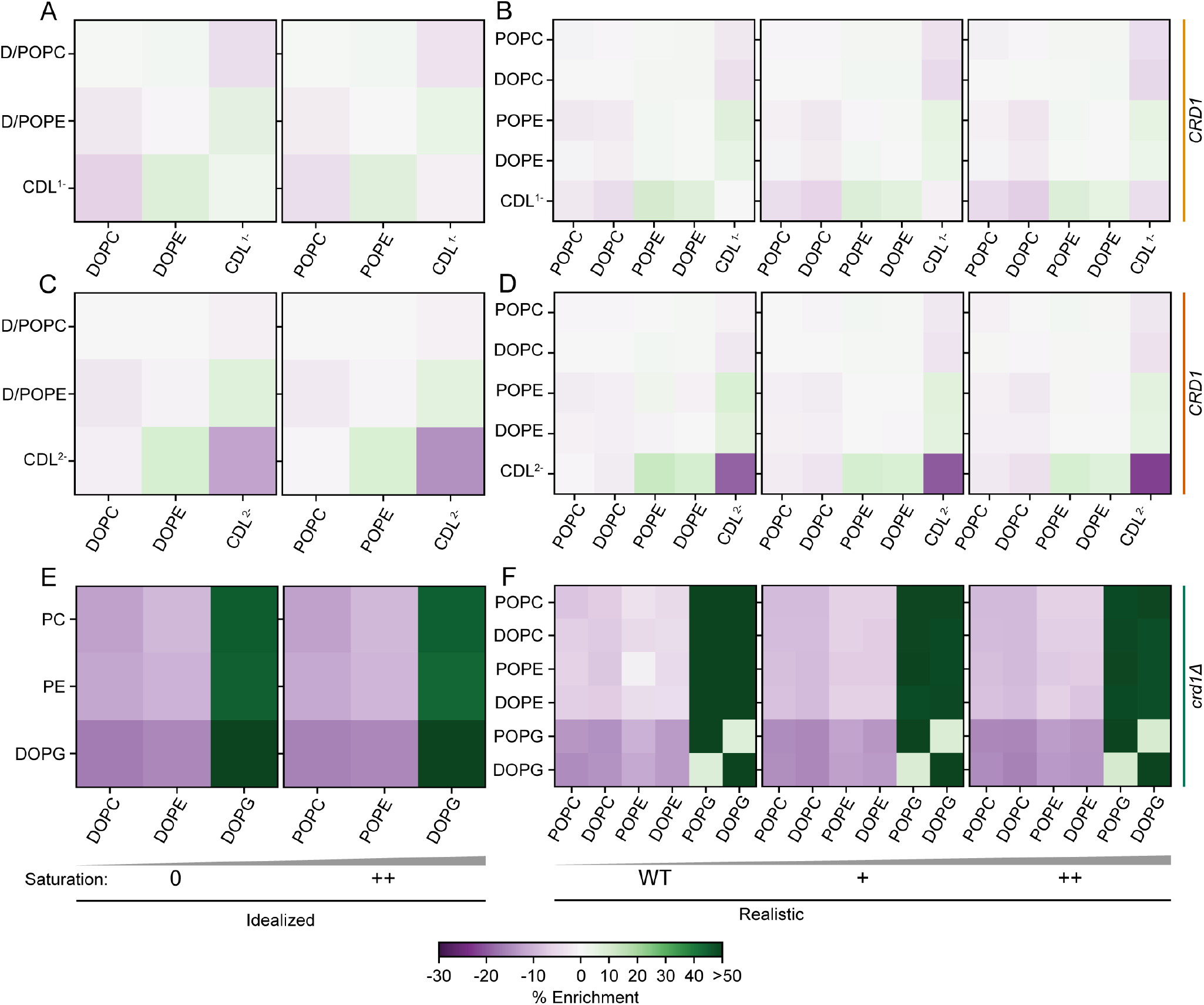
Quantification of the enrichment or depletion of y-axis lipid with x-axis lipid species interactions within a 1.5 nm radius compared to random. A, B) lipid enrichments of CDL^1 –^ containing idealized and realistic systems respectively. C, D) lipid enrichments of CDL^2 –^ containing idealized and realistic systems respectively. E, F) lipid enrichments of PG containing idealized and realistic systems respectively.

### Coupling between membrane undulations and lipid curvature

Having established that certain lipids have preferential lipid-lipid interactions, we next asked if there was any coupling between lipid clustering and transient fluctuations of the bilayer. Given that the bilayers we modeled with compositionally identical leaflets, we note that there are no net curvatures across the bilayer – just transient thermal undulations. Overlaying a rectangular grid onto the bilayer and interpolating a height field, we computed the mean curvature of the membrane at each grid point. Evaluating the time autocorrelation of the membrane mean curvature, we found that local curvatures become uncorrelated rapidly, within a few nanoseconds. The spatial autocorrelations were found to be local in nature and on the order of the grid spacing (i.e., 2 nm). We chose a timescale of 2.5 ns wherein the time autocorrelation of mean curvature drops to about 0.5 for all systems. Using this timescale, we computed a binned lipid density by binning the positions of lipids by type over 2.5 ns, Fig. 5A. To evaluate the extent of interactions between membrane undulations and each lipid, we computed the 2D spatial Pearson’s cross correlation coefficient of membrane mean curvature and lipid density, Fig. 5B. We found that in CDL containing systems, CDL and PE are negatively correlated with mean curvature while PC has a positive correlation. PG, on the other hand, negatively correlates with mean curvature to a lesser extent than CDL.

**Figure 5:**
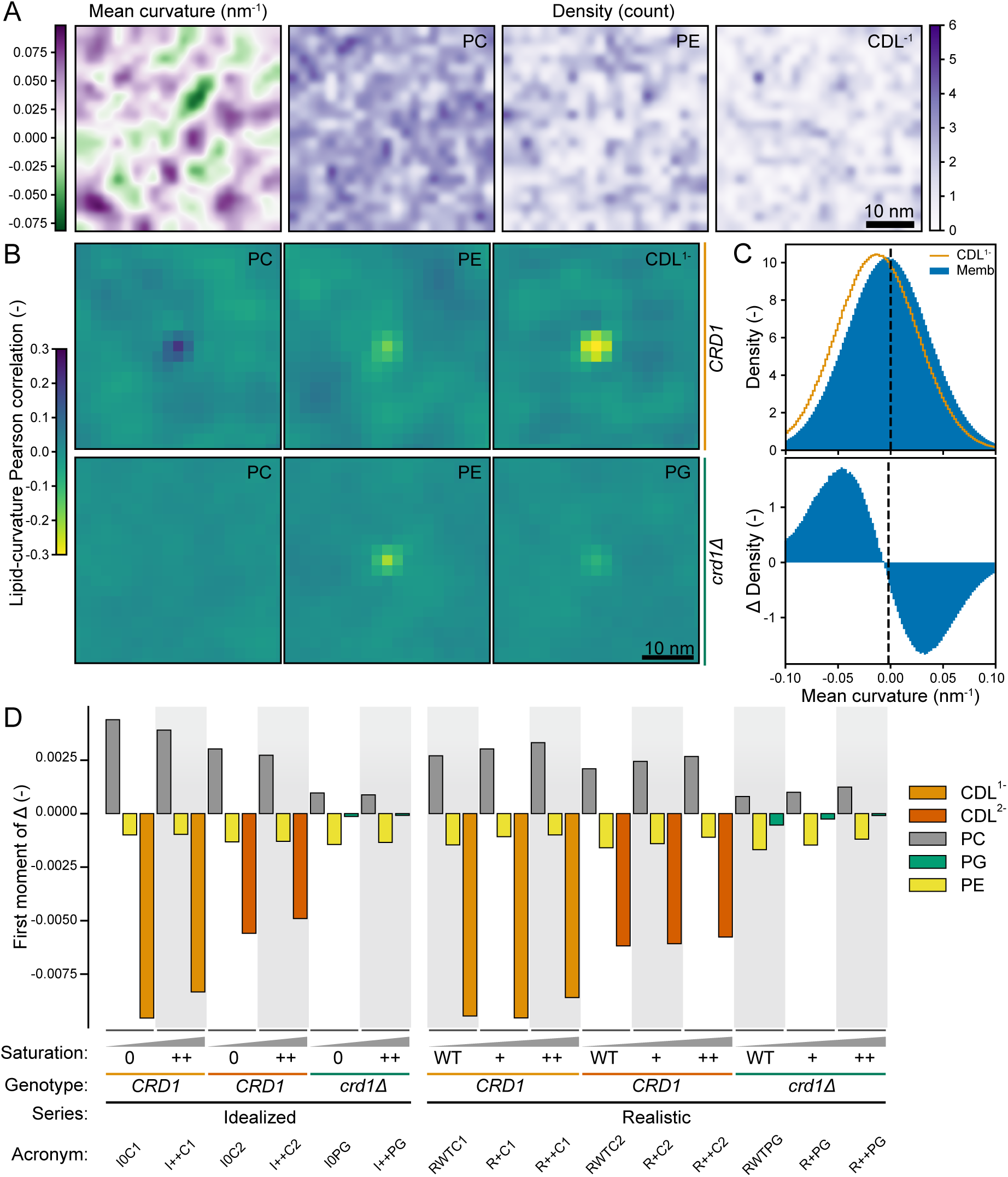
Correlations between membrane curvature and lipid localization. A) mean curvature of I0C1 with lipid densities (PC, PE, CDL^1 –^) obtained from averaging over 5 ns. Refer to Fig. S1 for the sign convention of the curvature. B) 2D spatial Pearson cross correlation coefficient of membrane mean curvature and lipid density for I0C1 and I0PG containing systems. C) Histogram of local curvature around CDL^1 –^ for IWTC1 system shown as orange line compared against all membrane curvatures shown in blue (top); (bottom) difference between the distribution of mean curvatures around a lipid of interest and the general membrane mean curvature distribution. D) quantification of the first moment of the difference between distribution of mean curvatures around a lipid of interest and the general membrane mean curvature distribution.

With the correlations between curvature and lipid density established, we further quantified the propensity of finding a particular lipid species at a given local curvature. We assigned each lipid a ‘curvature value’ by referencing the membrane curvature of the nearest grid point. We then compared the distribution of assigned curvature values for a given lipid type against the set of curvatures for all grid points to further quantify the curvature preference of each lipid type, Fig. 5C. The first moment of the difference between the lipid curvatures and the membrane curvature (ΔDensity) is a measure of the curvature weighted extent of deviation (i.e., differences at larger curvature values are weighted more heavily), Fig. 5D. Comparing the first moment of the difference, we found that CDL^1 –^ associates with negative mean curvatures to a greater extent than CDL^2 –^. PE was also found to be correlated with negative mean curvatures across all systems, albeit to a lesser extent than either CDL species. PC lipids are found to be enriched towards positive mean curvatures in all systems in a manner that appears, to an extent, relative to the shift towards negative mean curvatures of CDL and PE lipids. From these analyses, we conclude that CDL has the capability to localize to regions of negative mean curvature in a charge-dependent manner. We find that while PE also has a preference for negative mean curvature, CDL out competes PE for negative mean curvatures. Similarly, while PC has a limited preference for curvature, the presence of CDL partitioning can drive PC to other regions of the membrane.

### Cardiolipin inter-leaflet cross talk through membrane undulations

Having established that transient membrane undulations can drive localization of specific lipid types, we next considered whether there are curvature-driven effects between the two leaflets of the bilayer. Since the curvature of each leaflet is opposite relative to the other as constrained by the bilayer configuration, we next asked whether the curvature-induced aggregation in one leaflet affects the composition in the opposing leaflet. We computed the 2D spatial Pearson’s cross correlation of lipid densities across leaflets, Fig. 6A, and found that CDL is negatively correlated with itself across leaflets. That is local CDL enrichment/presence in one leaflet causes a depletion of CDL in the opposing region in the other leaflet. The other lipids, including PE despite the curvature preference, have weak correlations across leaflets. The Pearson’s correlation of unshifted lipid densities across leaflets is shown in Fig. 6B. The cross-leaflet depletion of CDL appears to be local in nature extending only ∼5 nm. Overall, we find that CDL is not only enriched at negative mean curvature undulations, but also depleted in the matching section of the opposing leaflets. This behavior is unique to CDL and not a detectable feature of other lipids like PE despite its large spontaneous curvature.

**Figure 6:**
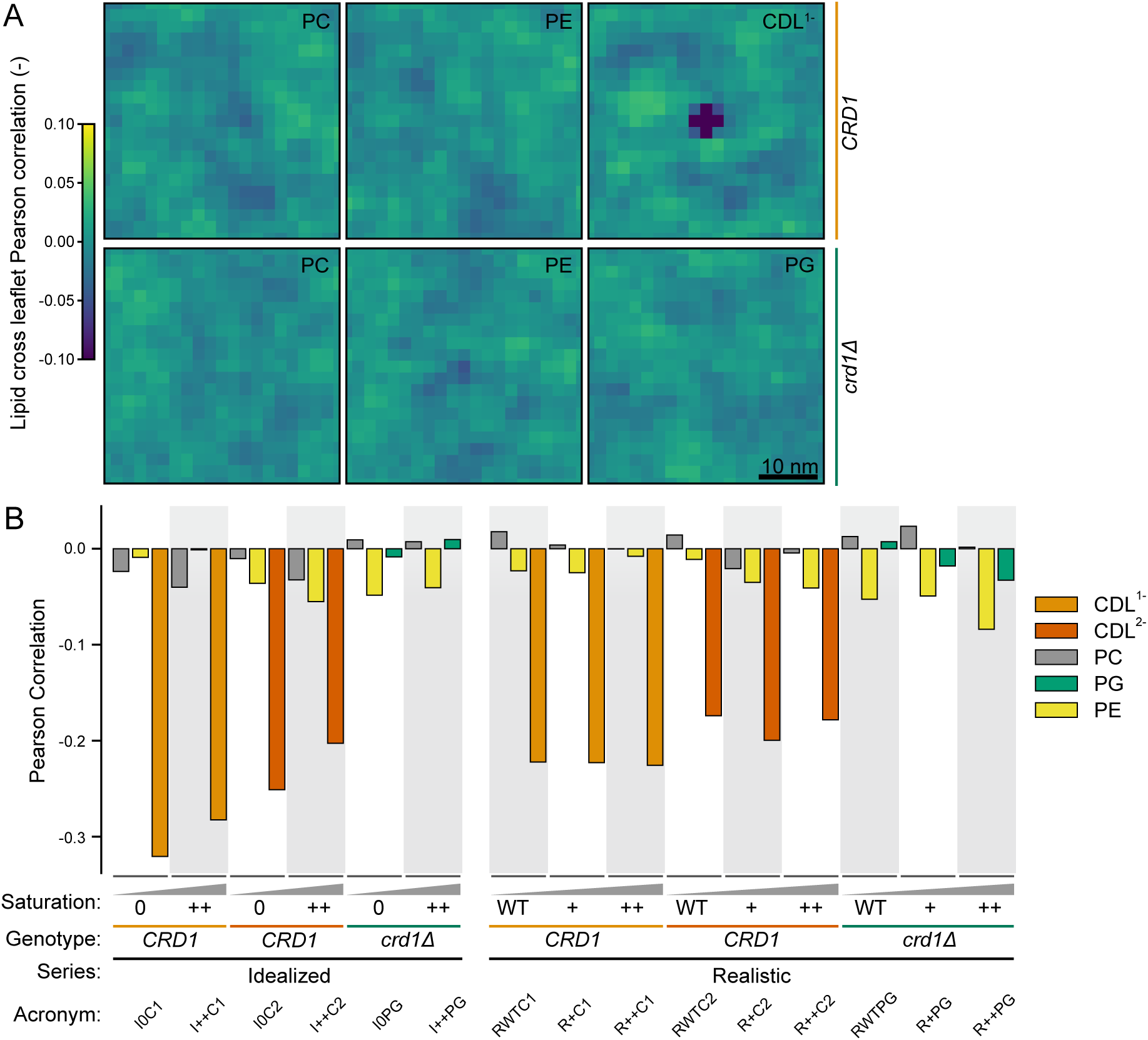
Pearson’s spatial cross-correlation of lipid densities across leaflets. A) Lipid-lipid cross-correlations for systems I0C1 (top) and I0PG (bottom). Lipid densities are generated by averaging over 5 ns. The reported cross-correlation is the average of 200 densities. B) Direct Pearson’s correlation of lipid densities across leaflets for all systems. This value corresponds to the central pixel in A, corresponding to the unshifted correlation between lipid density across leaflets.

## DISCUSSION AND CONCLUSIONS

In this work, we investigated the effects of changing lipid saturation on the material properties of the inner mitochondrial membrane. We found that increasing lipid saturation leads to an increase in bending rigidity and a general decrease in spontaneous curvature. Both trends point towards increasing difficulty in bending the membrane with increasing saturation. Continuum models of cristae formation suggest that the predicted parameter values lie in a range around a snap-through transition (i.e., where a small change in conditions leads to a drastic change in shape, like a jumping popper toy or slap bracelet) (9, 10, 48). The existence and proximity of the IMM to a snap-through instability can explain the experimental observation that there is a ‘breaking point’ of saturation where cristae morphology is lost. In our analysis, we identified unexpected spectral features in the power spectrum of membrane height undulations for CDL containing systems. These features were not predicted by the HC model and so we investigated whether they can be caused by heterogeneities in membrane organization.

With respect to the local lipid environment around lipids of a given species, we find that there are preferential interactions, which lead to enrichment or depletion of lipid partners. Such interactions have also been studied in the prior literature. Dahlberg and Maliniak [27] found that a decrease in charge led to an enhanced aggregation of CDL^1 –^ when compared to CDL^2 –^. They observed that CDL^1 –^ forms microdomains with negative curvature, but also that CDL^2 –^ does not. In contrast to this, we find that both CDL^1 –^ and CDL^2 –^ are transiently enriched at regions of negative mean curvature, to differing extents; CDL^1 –^ has a stronger correlation with negative mean curvature than CDL^2 –^. In apparent agreement with Ref. (27), we see that CDL^1 –^ is enriched around other CDL^1 –^, while CDL^2 –^ is depleted with respect to other CDL^2 –^. Thus, our simulations suggest that there appears to be a trade-off between specific interactions such as electrostatics and relaxation of bilayer stresses from geometric complementarity.

The correlation of other lipids with membrane curvature has also been observed in the context of asymmetric bilayers (49). Koldsø et al. [49] found that lipid species with different curvatures can partition to regions complementing their geometric preference. In our work, we modeled comparatively simpler compositions than Ref. (49), which do not undergo phase separation nor form more sophisticated heterogeneities. Nevertheless, even in our modeled symmetric bilayers, we observed transient lipid aggregation with membrane curvature. We found that lipids with strong curvature preferences such as CDL species can exclude other lipids, such as PC, from curved regions. These steric effects are composition dependent – PC shows no strong correlation with curvature in the *crd1Δ* systems (i.e., PG containing).

Other studies have also investigated the curvature partitioning of CDL and other lipids in systems with induced curvatures through buckling or other forcing mechanisms (30–32, 50–52). Our results showing that CDL has a greater propensity to associate with curvature than PE are in agreement with simulations of their partitioning in buckled membranes (29, 31, 53). Unlike models with forced curvature, our work identifies localization with curvature driven by thermal undulations even in flat, symmetric bilayers without external loading. While we calculated the cross-correlation between lipid density and membrane mean curvature, others have previously observed similar correlations using atomistic MD simulations of CDL correlated with *bilayer height* (43); The mean curvature is related through Eq. (21) to the height field.

In the case of CDL, we also observed negative correlations of self density across leaflets of the bilayer. Our observations are consistent with prior work showing differences in number density of local CDL across leaflets (27). Our work extends the observation, showing that the cross-leaflet effect is unique to CDL and not observed for the other lipid species which we modeled. We suggest that this is due to the strong curvature preference of CDL; a negative curvature in one leaflet corresponds to a positive curvature in the opposing leaflet. This is a possible mechanism to drive membrane heterogeneity even in symmetric membranes.

With respect to the discrepancies between experimental estimates of the stiffness and spontaneous curvatures of CDL containing bilayers: scattering experiments predict rigidities of ∼25 k_B_T and a reduced spontaneous curvature for CDL compared to PE (23–26); while other experiments investigating lipid partition to tubes pulled from a Giant Unilamellar Vesicle (GUV) predicted a spontaneous curvature of −1.1 nm^−1^ for CDL (21), a value similar to our estimate for CDL^2 –^, (−1.52 ± 0.02) nm^−1^ (cf. Fig. S7. Beltrán-Heredia et al. [21] suggest that lipid self-associative interactions drive this curvature localization despite the fact that no raft formation was observed in their work. Our simulations suggest that curvature complementarity may be the driving force leading to transient localization/’clustering’. In other words, if the diffusion of curvature-sensitive lipids is fast relative to the timescale of membrane height undulations, then the shape of the undulations will influence the localization of the lipids; The localization of CDL, which has a negative spontaneous curvature, to regions of membrane with matching curvature minimizes Eq. (1). This is a geometric mechanism which can be orthogonal to specific chemical self-interactions.

Atomistic MD simulations by others have also reported clustering to negative curvatures (43) and any curvature (34). In disagreement with our results, Wilson, Ramanathan, and Lopez [43] found that the membrane height fluctuations are not affected by CDL content (43). Estimated bending rigidities using real space fluctuations suggest a value of ∼30 k_B_T (33, 34), closer to scattering experiments. Perhaps the discrepancy is a result of the length scale difference between atomistic and coarse-grained simulations – systems may need to be sufficiently large to permit long wavelength undulation modes to support partitioning and thus curvature softening (35). From experiments, it has been suggested that changes to local membrane bending rigidity (from changing lipid composition due to demixing) can lead to curvature sorting (54). In line with our predicted rigidities, larger length scale experimental measurements using micropipette aspiration of mitochondrial lipid extracts estimate a bending rigidity of 5 k_B_T (22). Overall, we suggest that geometric frustration of CDL in planar membranes facilitates the development of curvature-undulation reinforcing heterogeneities, which appear as an effective softening of the membrane (35, 55). It is possible that the discrepancy between PE and CDL spontaneous curvature and rigidity ordering between experiments can be explained by difference in the length- and time-scale of experimental set up and fitting approach. With respect to spontaneous curvature estimates, measurements in giant vesicles via flicker spectroscopy or micropipette aspiration will capture the contributions of compositional heterogeneities driven by curvature coupling while such heterogeneities may only occur in the planar orientation of H_II_ systems and this may not be captured by SAXS. Regarding bending rigidity, the timescale of measurements may lead to the observed differences. Micropipette aspiration and flicker spectroscopy report equilibrium values subject to the influence of compositional heterogeneity while other approaches which infer rigidity from fast relaxing processes will not reflect the diffusional softening (35).

Beyond the membrane, the change in lipid chemistry may influence their interactions with embedded proteins. The preferences of lipids to interact with certain proteins in a structure dependent manner has been called lipid fingerprints (56, 57). The principles of how the lipid bilayer organizes influences its properties as a solvent for these proteins. CDL is asymmetrically distributed in the IMM. Curvature coupling of CDL within and across leaflets may aid in the localization of transmembrane and integral proteins on the opposing leaflet. Our study suggests that lipid spontaneous curvature can drive heterogeneities in the bilayer adding to the emerging functional role that lipid composition plays in organelle physiology (58).

## AUTHOR CONTRIBUTIONS

CTL, KV, IB, and PR designed research. CTL performed research with feedback from KV, IB, and PR. CTL ran the simulations, wrote the scripts to execute analysis, and analyzed the data. CTL wrote the original draft and CTL, KV, IB, PR reviewed and edited the manuscript.

## ACKNOWLEDGMENTS

The authors would like to thank Edward Lyman who provided helpful discussions. We also thank Honor Akenuwa and Nate Linden-Santangeli for critical review. This work was supported by grants from the Office of Naval Research (ONR N00014-20-1-2469 to PR), a Moore-Simons Project on the Origin of the Eukaryotic Cell (GBMF-9734 to IB, PR), and National Institutes of Health (R35-GM142960 to IB). KV was supported by the NIH Molecular Biophysics Training Grant (T32-GM008326). CTL was supported in part by a Kavli Institute for Brain and Mind Postdoctoral Fellowship. Molecular dynamics simulations were run on hardware hosted by the Triton Shared Computing Cluster (59).

## A SUPPLEMENTARY FIGURES

**Figure S1:**
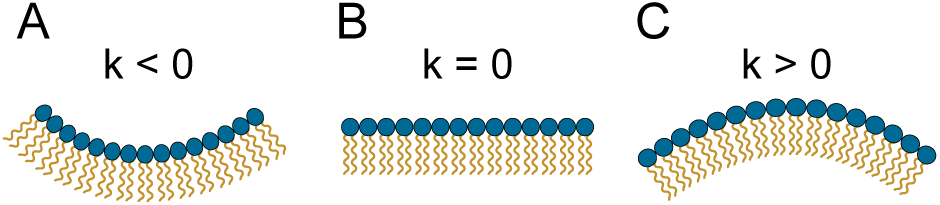
Choice of curvature convention used in this work. Blue dots correspond to headgroups and orange squiggles to the acyl tails.

**Figure S2:**
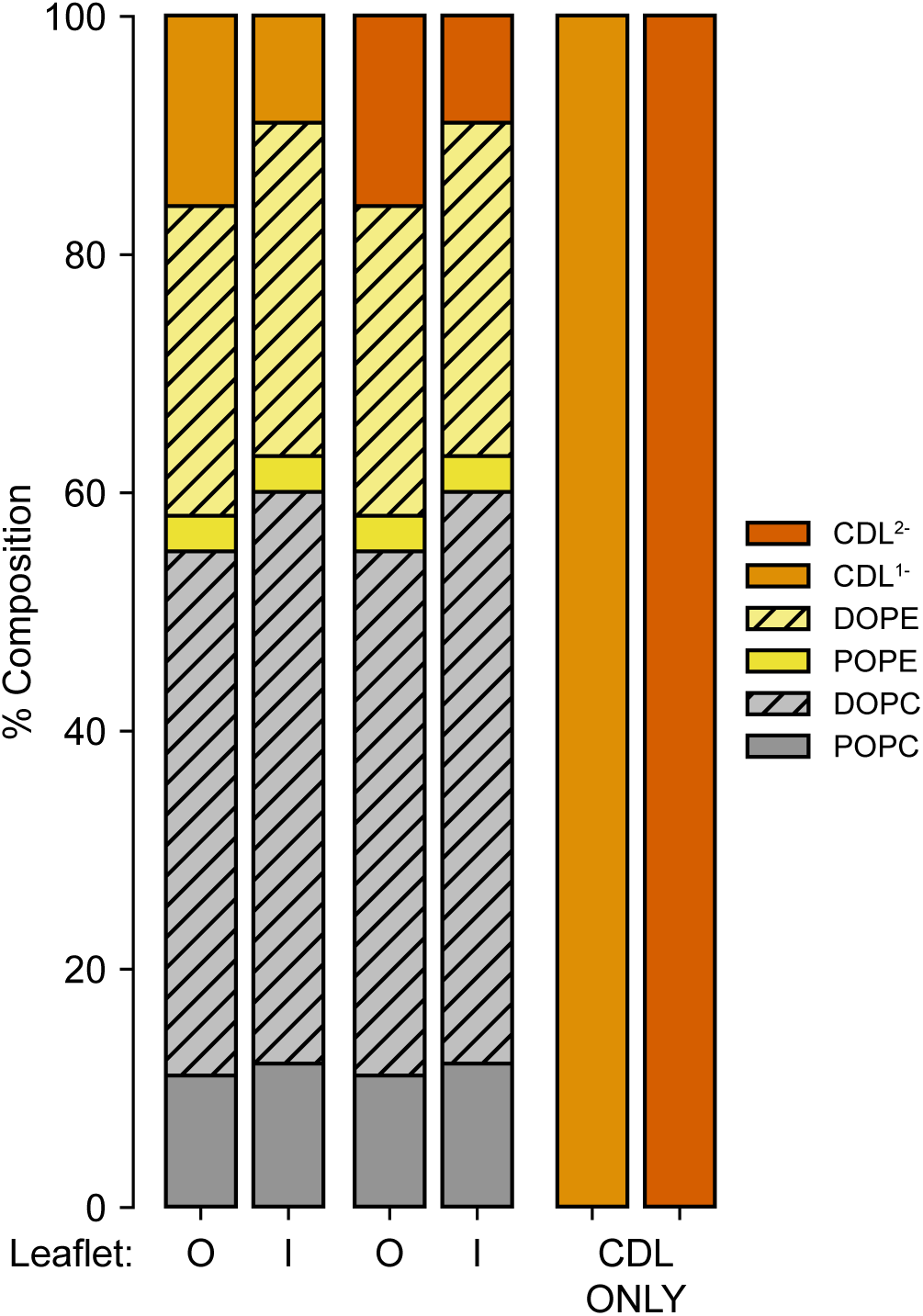
Compositions of other systems modeled including symmetric outer- and inner-leaflet like compositions of the IMM and pure CDL^1 –^ and CDL^2 –^ containing systems.

## SUPPLEMENTARY MATERIAL

An online supplement to this article can be found by visiting BJ Online at http://www.biophysj.org.

## B METHODS

### B.1 Molecular dynamics

All codes for simulation set up and analysis can be found online (69). Software versions used are documented in Table S2.

From configurations generated by insane.py, systems are minimized first using soft-core minimization of both Van der Waals and charge interactions steepest-descent for 1000 steps followed by conventional steepest descent minimization for an additional 1000 steps. Minimized systems then undergo restrained equilibration where harmonic restraints holding the lipid head groups in place are relaxed from 200–50 kJ mol^−1^ nm^−2^ with concomitant increase of timestep from 2–10 fs over a series of steps. All equilibration steps are performed in the NPT ensemble, using the Berendsen semiisotropic barostat (70).

**Table S1:**
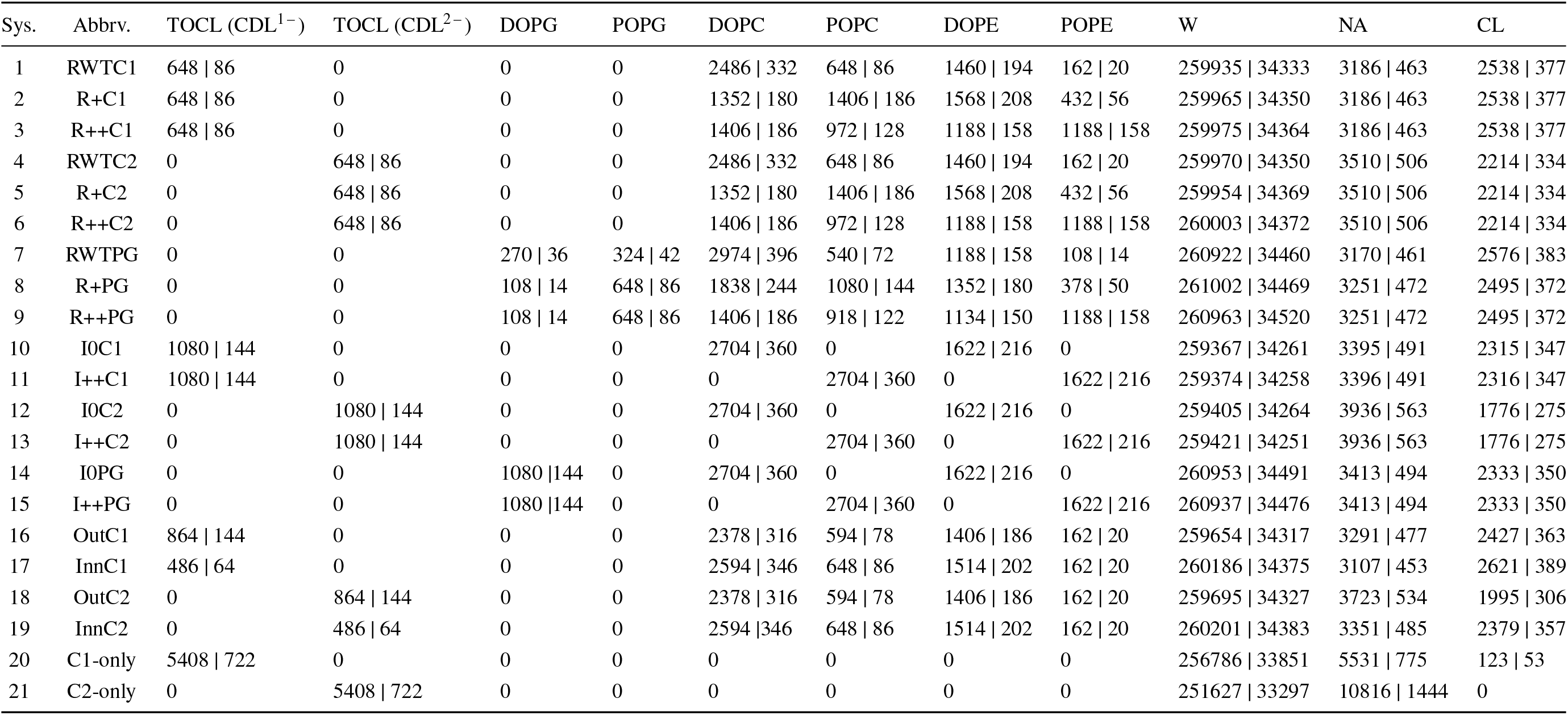
Compositions for each system (large | small) with abbreviations. Abbreviation nomenclature follows corresponding to idealized/realistic (I, R), wild type/increased saturation (WT/0, +,++), and CDL^1 –^ /CDL^2 –^ /PG (C1, C2, PG).

**Figure S3:**
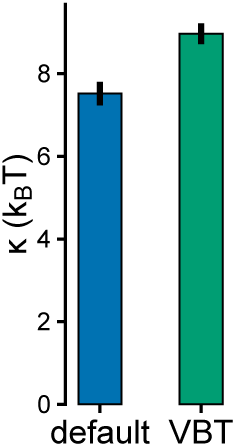
Comparison of the estimated bending modulus for the RWTC1 composition with simulation control parameters in this study (default) and following the recommendations by Kim, Fábián, and Hummer [45] (VBT). In agreement with their results suggesting that the default simulation settings can miss interactions leading to increased membrane crumpling and height fluctuations, we find that the default settings underestimate the bending modulus by ∼20 % (i.e., the systems appear to be easier to bend owing to the buckling).

**Figure S4:**
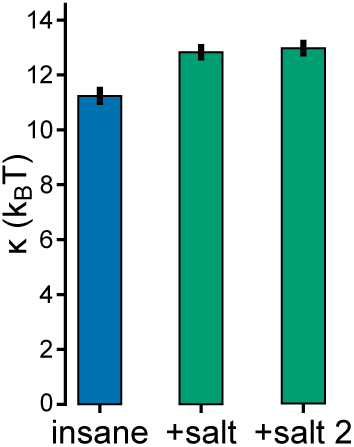
Comparison of the estimated bending modulus for the I++C2 system with ions added by insane.py which adds salt followed by removal of ions to neutralize the system (insane) or neutralizing net charge followed by adding salt (+salt, +salt 2; two replicates with identical conditions). The latter approach leads to more total ions in the system with the number of paired ions corresponding to the target salt concentration. The system with additional ions (+salt) have an increase in bending rigidity compared to systems generated by insane.py (insane). This may be due to the screening of CDL^2 –^ -CDL^2 –^ interactions by the extra ions leading to reduced transient aggregation and a corresponding reduction of curvature softening.

**Table S2:** Major software and versions used in this study.

| Software | Version (Zenodo ref.) | Refs. |
| --- | --- | --- |
| Gromacs | 2022.1 or 2024.4 (60) | (61) |
| MDAnalysis | 2.2.0 | (62) |
| numpy | 1.21.5 | (63) |
| Scipy | 1.7.3 | (64) |
| matplotlib | 3.6.3 (65) | (66) |
| membrane-spectral-analysis | 0.1.0 (67) | – |
| insane.py | 1.1.dev0 | (37) |
| VMD | 1.9.4a55 | (68) |

**Figure S5:**
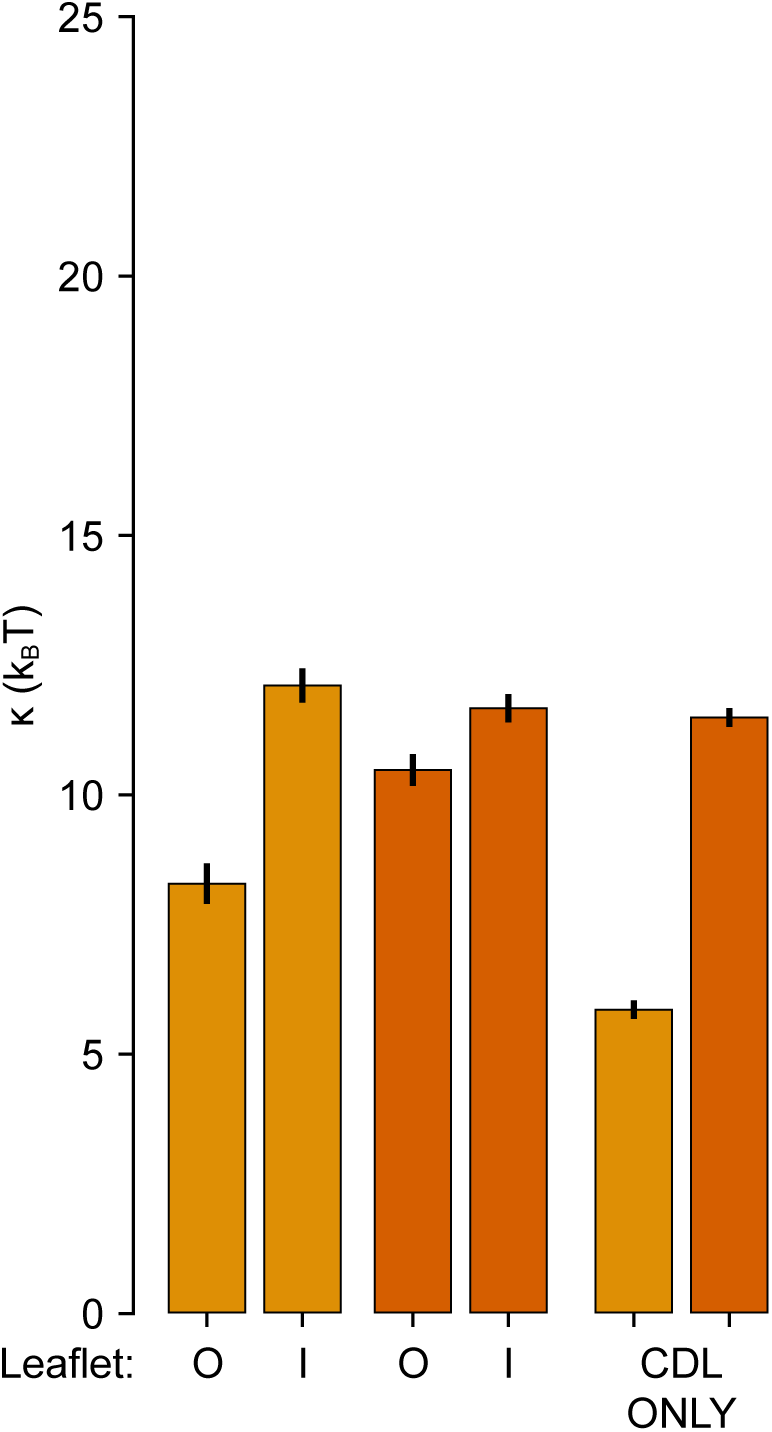
Estimated bending modulus for symmetric systems with outer leaflet-like (O) and inner leaflet-like (I) of the of the IMM composition and cardiolipin only.

**Figure S6:**
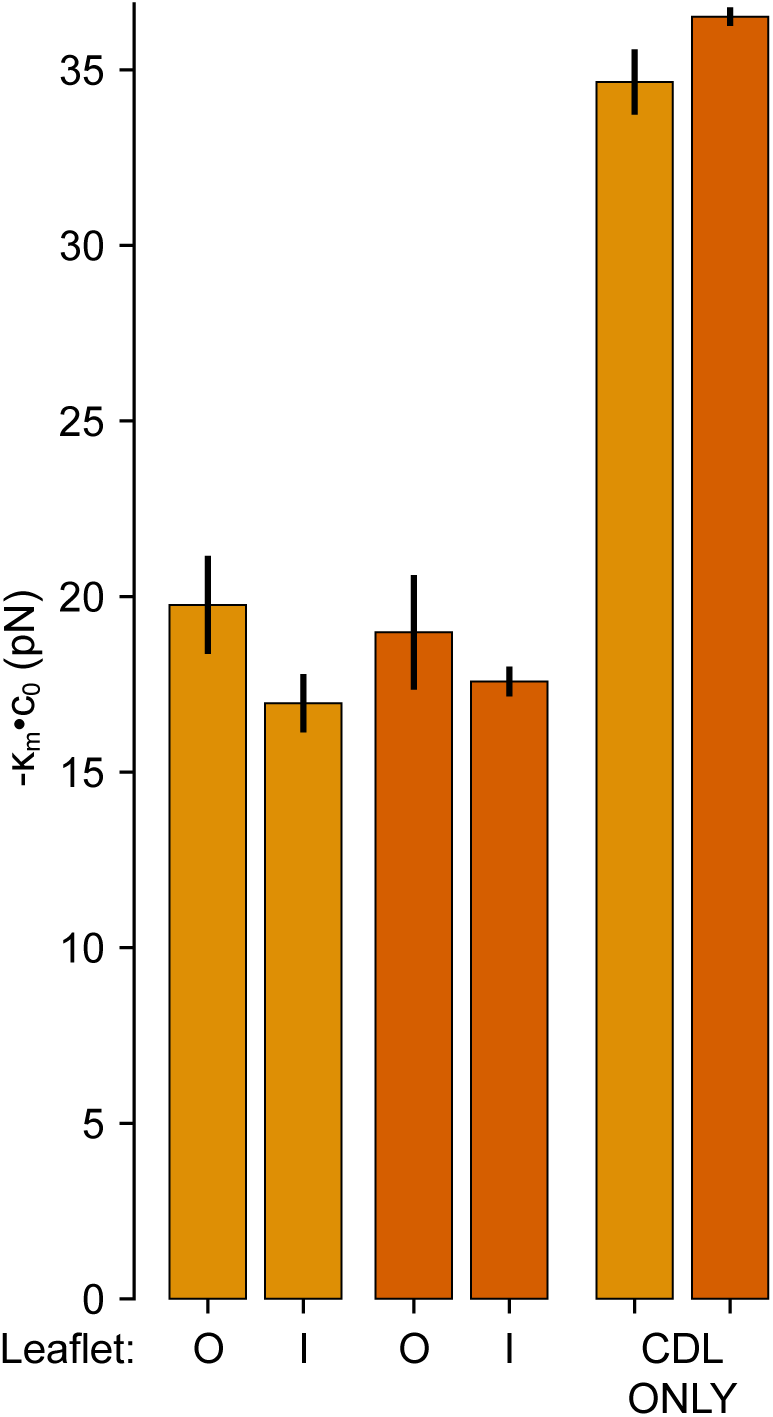
Estimated bending moment for symmetric systems with outer leaflet-like (O) and inner leaflet-like (I) of the IMM composition and cardiolipin only.

**Figure S7:**
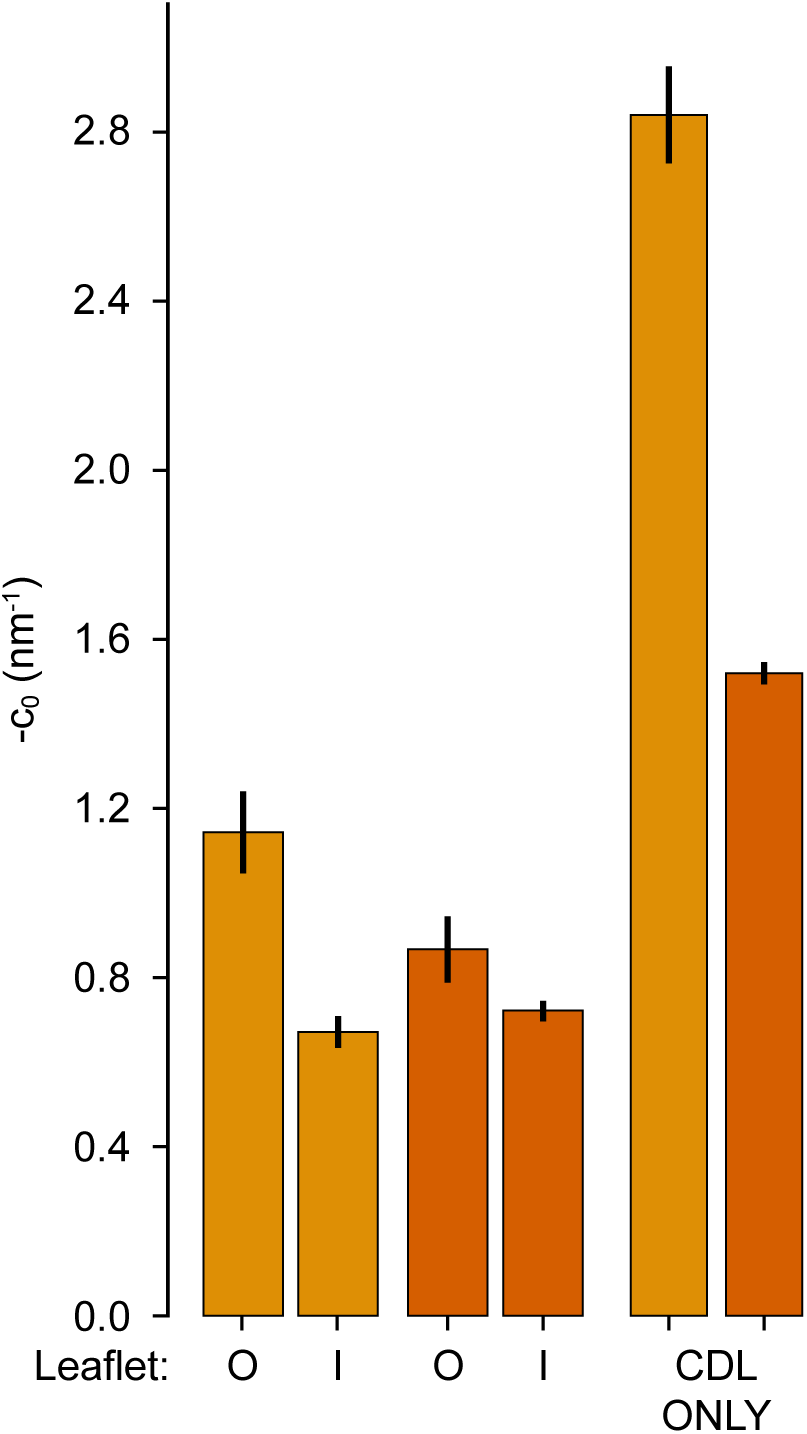
Estimated spontaneous curvature for symmetric systems with outer leaflet-like (O) and inner leaflet-like (I) of the IMM composition and cardiolipin only.

#### B.1.1 Estimation of bending modulus

We start by summarizing the approach to derive the well-known expression relating the bending modulus and the height fluctuation spectrum (15, 19, 44, 46, 71). Starting from the Helfrich-Canham Hamiltonian,

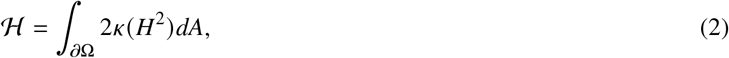

where *ℋ* is the energy of the system, *a*n is the surface of the domain n, *K* is the bending modulus, and *H* = (*k*_1_ + *k*_2_)/ 2 is the mean curvature of the surface. We note that Eq. (2) deviates from Eq. (1) by the spontaneous curvature term. Since the modeled membrane are up-down symmetric in composition, we expect that there should be no net membrane spontaneous curvature.

Assuming the membrane shape is an approximately flat patch with small deviations from a plane, we can express the energy in the linear Monge gauge,

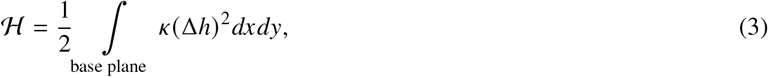

where Δ is the Laplace operator, and *h* : (*x, y*) ↦ *h*(*x, y*) is the height. Note in the previous step that we have used a small gradient approximation, 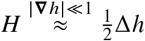.

From the linear Monge gauge, the shape of the system can also be represented in Fourier space. Assuming that the membrane shape is defined on an *L* × *L* plane with periodic boundary conditions, we get the Fourier pair,

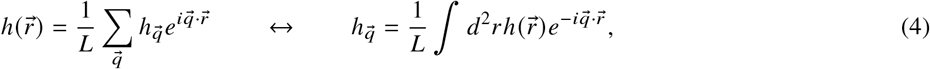

where 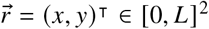, and

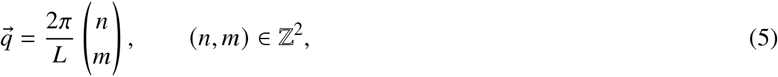

is the wave vector.

Evaluating the gradients of *h* in Fourier space and substituting, we can re-express the Hamiltonian in a Fourier basis,

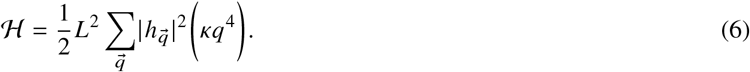

Noting that the wave vectors are orthogonal and harmonic, we can apply the equipartition theorem to get

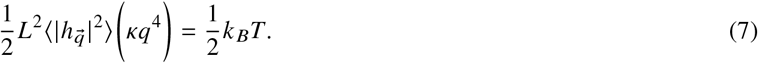

Rearranging, we get

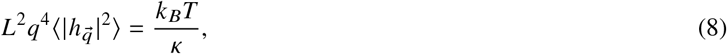

where *k* _*B*_*T* is the product of the Boltzmann constant and temperature. We will fit the height fluctuation spectra to Eq. (8) to estimate the bending modulus.

#### B.1.2 Statistical analysis of fluctuation spectra

The trajectories were analyzed using MDAnalysis. We note that algorithmically, we do not define a continuous height function for the membrane. Instead, we derived a discrete height field on a regular grid by sampling a piecewise cubic interpolation of a tessellated point cloud made up of membrane surface beads (Scipy griddata). We used the PO4 and GL0 beads, both present in the head group region of the various lipids, to represent the surface of the bilayer. Lipids corresponding to the upper and lower leaflets were identified using a graph clustering algorithm, LeafletFinder, available from MDAnalysis. A parameterization, *h* : (*i, j*) → *z*, of each leaflet on a regular quadrilateral mesh with 2 nm was obtained by tessellating the point cloud of a given leaflet in the *x* and *y* directions followed by 2D cubic interpolation. The tessellation is performed in the plane of the bilayer using information from the periodic boundaries to produce a large 3 × 3 system. In this manner, the height field at the boundaries of the central image are representative of the membrane surface and void of boundary artifacts. The continuous Fourier transform on the right hand side of Eq. (4) is approximated by the discrete Fourier transform,

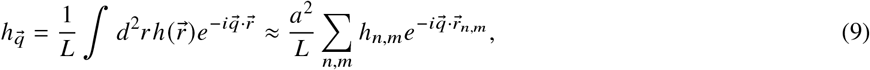

where *a* is a uniform grid spacing. The discrete 2D Fourier transform was calculated using utilities from Scipy and the resulting 2D height fluctuation spectra was converted to a 1D spectra by radial averaging (cf., Ref. (44)). Altogether, we fit a constant to the following expression,

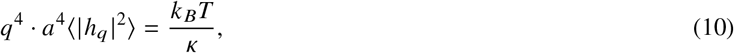

to estimate the bending modulus.

For uncertainty quantification we followed the approach outlined by Ergüder and Deserno [46]. Noting that each of the undulatory modes in Fourier basis are orthogonal and thus independent, we can consider the evolution of the power within each undulatory mode as an independent dynamical variable. To obtain an estimate of the uncertainty we must consider if the sampled data points are independent and identically distributed. Since energy within each undulatory mode is owing to thermal fluctuations, for a well thermalized system, we expect the process to be stationary. Meanwhile, since the membrane conformations are sampled at some arbitrary write out frequency, there may be correlations in time.

To obtain an unbiased variance of correlated data, one approach is to calculate the statistical inefficiency, *s*, which is given by (72–74),

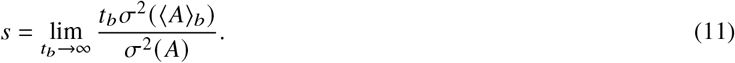

Using this statistical inefficiency, we can correct the variance to obtain an unbiased estimate,

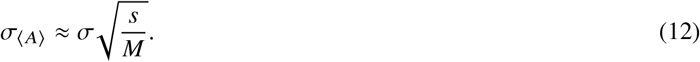

A traditional approach would be to apply this style of block averaging to a sequence of estimated and possibly correlated bending rigidities from segments of the simulation to get an estimate of the uncertainty. The limitation of this approach is that the underlying relationship between the bending rigidity and amplitude of height fluctuations is only true on average. When subdividing the trajectory into blocks, at even modest block lengths we found that there is insufficient data such that thermal noise is not averaged out and dominates the error estimate.

Instead, by treating the power per wave mode as a dynamical process in and of itself, we can perform block averaging of each components of the power spectrum. The variance of power within each undulatory mode, can be estimated by fitting to the following relation (46),

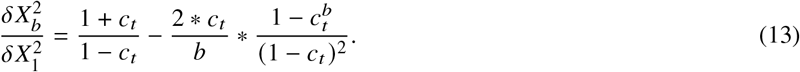

**Figure S8:**
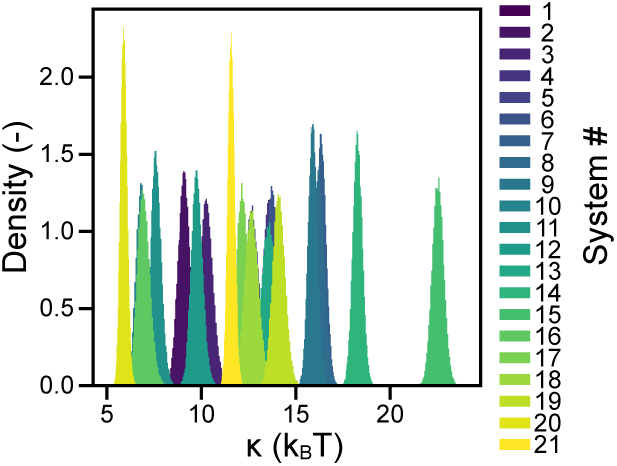
Distribution of bending moduli values for all modeled system from parametric bootstrapping.

Given the central theorem of statistics, the distribution of power values for each mode is Gaussian and thus we can use parametric bootstrapping to generate new datasets. By fitting bending modulii to many sets of bootstrapped data, we obtained a boostrapped distribution, shown in Fig. S8 from which we determined the mean and standard deviations.

#### B.1.3 Calculating the spontaneous curvature from the stress profile

**Figure S9:**
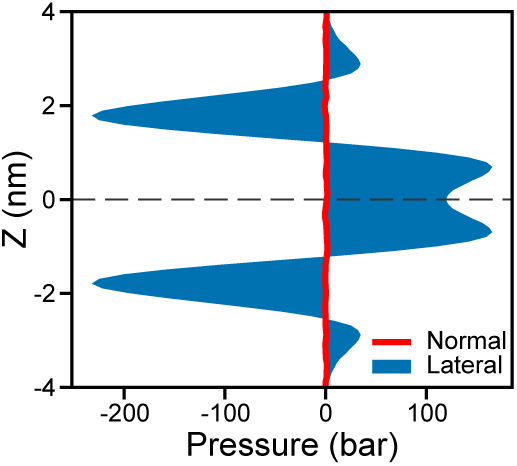
Example lateral and normal pressure profiles from the I++PG condition. The lateral pressure profile is given by Eq. (14).

Lateral pressure profiles are calculated by using MDstress software (75), gmx-ls. To analyze the system, we ran an extra 200 ns analysis trajectory after the 10 µs production on a fine write out rate of 5 ps, as recommended by Vanegas, Torres-Sánchez, and Arroyo [75].

We then centered the bilayer in the simulation box, and calculated the stresses on a 0.1 nm grid. Gmx-ls calculates the stress tensor of the system, from which the lateral pressure profile can be derived. The lateral pressure, *π*(*z*), can be calculated as the difference between the normal stress tensor and the average of the x, y plane tensors, *σ*,

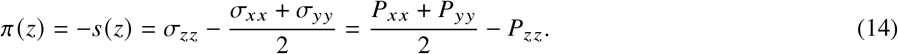

An example lateral and normal pressure profile for the I++PG condition is shown in Fig. S9.

The first moment of the lateral pressure profile corresponds to the bending moment (16, 17, 76–81),

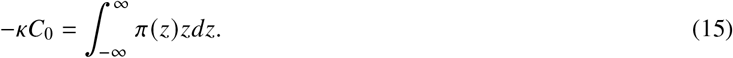

To compute the monolayer bending moment we integrate over each leaflet,

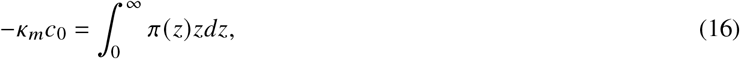

while further assuming that the monolayer bending rigidity is half of that of the bilayer, *K*_*m*_ = *K* /2.

The zeroth and first moments are evaluated by integrating a cubic interpolation of the lateral pressure profile for each system using Scipy (64). Error for the lateral pressure calculation is computed by splitting each 200 ns trajectory into three non-overlapping parts and evaluating the conventional mean and standard deviation.

#### B.1.4 Calculation of neighbor propensity

The local compositional enrichment of lipids was investigated by counting the lipids found within a range around a given lipid type. We define the location of each lipid as the coordinate of either the PO4 or GL0 bead in the head-group region. We compute a neighbor count, *C*,

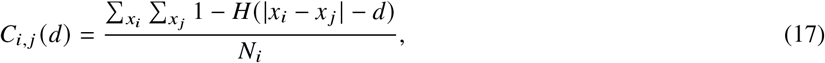

which corresponds to the mean number of lipids of type *j* around any lipid of type *i*; *H* is the Heaviside function, | *x*_*i*_ −*x* _*j*_| is the distance between lipids *x*_*i*_ and *x* _*j*_, and *N*_*i*_ is the number of lipids of type *i*. We compute *C* for each frame of the trajectory and take the mean 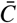 to represent the averaged lipid counts around a query lipid. The empirical probability *P* of finding a lipid *j* around lipid *i* is then given by,

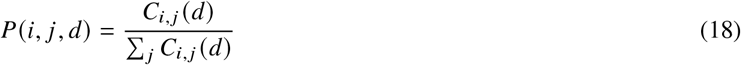

This empirical probability can be compared against the naïve probability which assumes a random distribution which implies that the probability of finding a lipid *j* around lipid *i* is,

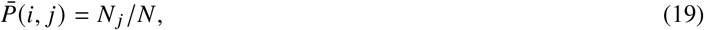

where *N* _*j*_ is the number of lipid type *j* and *N* the total number of lipids. The enrichment, *E*, in comparison against a random lipid distribution is given by,

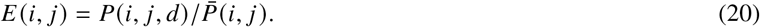

#### B.1.5 Estimation of local curvature, lipid densities, and correlations

The mean curvature of the membrane was computed using a modified version of the Membrane-Curvature library. We aim to parameterize a surface given by the location of P04, P041, P042, and GL0 beads which correspond to the head group of the lipids. Lipids corresponding to the upper and lower leaflets were identified using a graph clustering algorithm, LeafletFinder, available from MDAnalysis. The geometry of the each leaflet, represented by a regular quadrilateral mesh with 2 nm spacing, was computed by tessellating the point cloud of a given leaflet in the *x* and *y* directions. We performed a 2D cubic interpolation of the tessellated point cloud to produce a parametrization of the surface, *S* : (*i, j*) → *z*. The tessellation prior to fitting helps to prevent boundary artifacts in the interpolated surface geometry. The mean curvature, *H*, at a given point of the quadrilateral mesh can then be computed by,

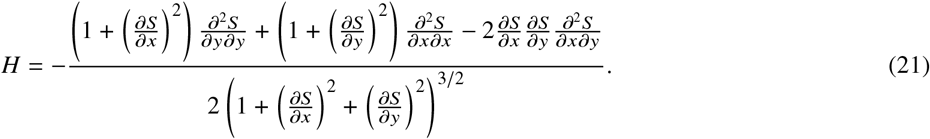

**Figure S10:**
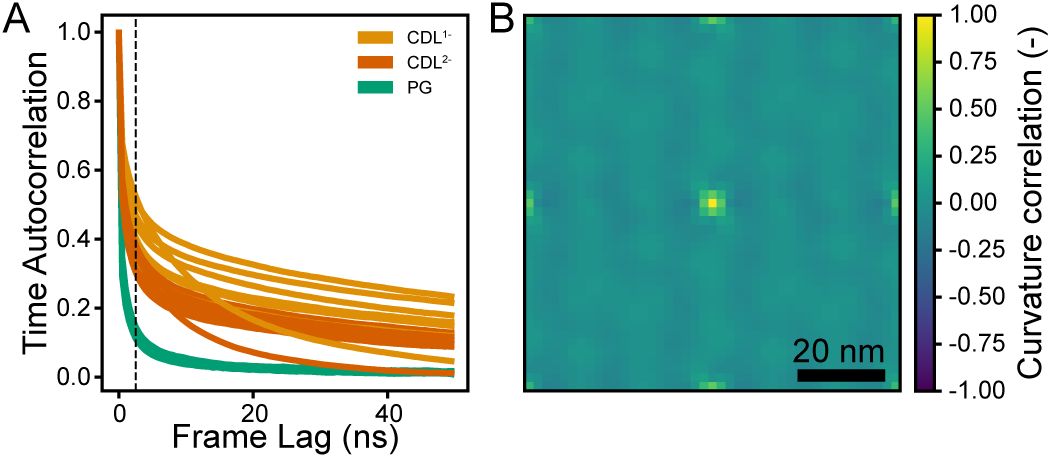
Curvature autocorrelation functions in time for all systems (A) and a representative system, I0C1, in space (B). Other systems have similar spatial curvature autocorrelations to the representative system. Dotted line in A indicates the threshold chosen for averaging frames to produce representative lipid densities, membrane curvatures, and correlations shown in Fig. 5.

The time autocorrelation of the mean curvature for a given point (*i, j*) is given by,

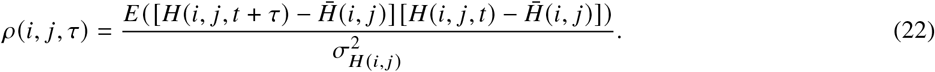

The mean time autocorrelation averaged all grid points is shown in Fig. S10A. We choose a timescale of 5 ns which corresponds to a period of time over which a given point has memory of it’s past mean curvature. This timescale will be used in later analysis for calculation of lipid densities.

The spatial autocorrelation of the mean curvature for a given frame at time *t* is given by,

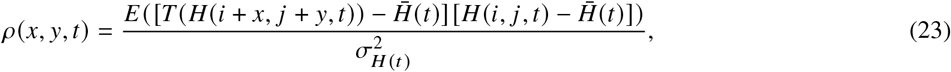

where *T* (·) is a wrapping transform to account for periodic boundary conditions. The mean spatial autocorrelation which averages over all frames is shown in Fig. S10B.

In order to compare the local lipids with the membrane curvature, we perform two analysis: 1) assignment of a local curvature to each lipid followed by binning; and 2) cross correlation of lipid densities and membrane curvatures smoothed over time.

We assign a curvature value to each lipid by choosing the mean curvature of the closest grid point. Special care was taken to consider periodic boundary conditions. Furthermore, since the MD simulations are run in the NPT ensemble, the grid corresponds the frame with the smallest XY dimension across the trajectory, we filter out lipids too far from interpolated grid. A histogram of curvatures experienced by each lipid was built by binning the assigned lipid curvature values differentiated by lipid type over the trajectory.

To compute the lipid density, we count the lipids associated with each grid point over the aforementioned 5 ns timescale, and normalize such that the sum over the lipid density is unitary. The 2D Pearson spatial cross-correlation for lipid density with the mean curvature, is computed using Eq. (23) with the appropriate modifications. Similarly the 2D Pearson spatial cross correlation of lipids across leaflets is evaluated.

